# Pyruvate dehydrogenase kinase expression profile is a biomarker for cancer sensitivity to dichloroacetate-mediated growth inhibition

**DOI:** 10.1101/2023.09.18.557101

**Authors:** Bevan P Gang, Melissa Rooke, Ramon C Sun, Samagya Banskota, Sheenu Mishra, Jane E Dahlstrom, Anneke C Blackburn

## Abstract

**Background:** Cancer cells favour glycolysis and lactate production over mitochondrial metabolism despite the presence of oxygen (the Warburg effect). Increased pyruvate dehydrogenase kinase (PDK) activity contributes to this glycolytic phenotype. Dichloroacetate (DCA) is a PDK inhibitor with anti-cancer potential that inhibits all four isoforms of PDK but with differing potencies, thus expression of different isoforms may determine sensitivity to DCA.

**Methods:** The association of sensitivity to growth inhibition by DCA, on-target effects of DCA and expression of all four isoforms of PDKs in a range of epithelial cancer cell lines was investigated *in vitro* and *in vivo*.

**Results:** DCA inhibited growth of cancer cells *in vivo* and *in vitro,* reduced pyruvate dehydrogenase phosphorylation and reduced lactate production. The magnitude of the effect of DCA on growth was variable and correlated with the PDK expression profiles of the cells, with low expression of PDK3 (highest K_i_ for DCA) conferring the highest sensitivity towards DCA. PDK2 siRNA-knockdown inhibited growth to a similar extent to DCA, whilst PDK3 knockdown significantly increased sensitivity to DCA.

**Conclusion:** The PDK expression profile is a potential biomarker for sensitivity to DCA, and should be considered when translating PDK inhibitors into clinical use.

## Introduction

Deregulated metabolism as a phenotype of cancer cells has recently received an immense amount of attention, and is considered an emerging hallmark of cancer (Cairns *et al*, 2011; Hanahan & Weinberg, 2011; Possemato *et al*, 2011; Sasaki *et al*, 2012; Zaugg *et al*, 2011). In the 1920s, Otto Warburg discovered metabolic deregulation in cancer cells from his observation that cancer cells prefer to use glycolysis even in the presence of oxygen (Warburg, 1956). The Warburg effect results in the diversion of pyruvate metabolism from mitochondrial oxidation to lactate. The fate of pyruvate is determined by the balance between the activity of lactate dehydrogenase and pyruvate dehydrogenase (PDH). Thus PDH determines the ratio of coupling between glycolysis and mitochondrial oxidation. There are four pyruvate dehydrogenase kinase (PDK) isoforms, PDK1-4, which inhibit pyruvate dehydrogenase (PDH) activity by phosphorylating three serine residues (Ser232, 293 and 300) on each of the two E1α subunits of the enzyme (Kolobova *et al*, 2001; Korotchkina & Patel, 2001; Papandreou *et al*, 2011). PDK1 is expressed predominantly in heart tissue, PDK2 is ubiquitous, PDK3 is expressed in testis, and PDK4 is expressed in heart and muscle (Bowker-Kinley *et al*, 1998). HIF1 is a transcription factor composed of HIF1α (stabilized in low oxygen) and HIF1β (constitutive) subunits. HIF1 is responsible for the major gene expression changes in response to hypoxia, including induction of PDK1 and PDK3 expression (Lu *et al*, 2008; Papandreou *et al*, 2006). PDK1 activity can also be increased by phosphorylation by oncogenic tyrosine kinases such as FGFR1 and BCR-ABL (Hitosugi *et al*, 2011) and expression can be increased via HIF1 interaction with oncogene MYC (Kim *et al*, 2007). Thus the PDK-PDH axis can integrate signals from growth factors, oncogenes and the microenvironment and is a central regulatory hub for tumor metabolism.

Inhibition of PDKs in cancer cells reverses the Warburg effect by activating PDH and redirecting pyruvate metabolism back into the mitochondria (Bonnet *et al*, 2007; McFate *et al*, 2008). PDK1 has been the predominant focus of previous studies (Cairns *et al*, 2007; Hitosugi *et al*, 2011; Kim *et al*, 2007; McFate *et al*, 2008) whilst the contribution of the other three PDKs in tumor metabolism has been largely ignored. Dichloroacetate (DCA) is a small-molecule PDK inhibitor that has been used in the clinic for lactic acidosis disorders for over three decades (Stacpoole & Greene, 1992; Stacpoole *et al*, 1983). DCA has been demonstrated to halt tumor growth *in vivo* (Bonnet *et al*, 2007; Chen *et al*, 2009; Sanchez-Arago *et al*, 2010; Sun *et al*, 2010), and in glioblastoma patients could normalize the mitochondria, and induce apoptosis in tumors (Michelakis *et al*, 2010). However, the effect of DCA on growth inhibition has been variable *in vitro* and *in vivo* in several cancer types (Anderson *et al*, 2009; Cairns *et al*, 2009; Cairns *et al*, 2007; Cao *et al*, 2008; Chen *et al*, 2009; Sanchez-Arago *et al*, 2010; Sun *et al*, 2010; Sun *et al*, 2009; Wong *et al*, 2008), and may require concentrations >10 mM to induce apoptosis at (Heshe *et al*, 2010; Madhok *et al*, 2010; Stockwin *et al*, 2010). The four PDKs vary in their sensitivity towards DCA inhibition. PDK2 is most sensitive (K_i_ 0.2 mM), followed by PDK4 (K_i_ 0.5 mM), PDK1 (K_i_ 1 mM) and lastly PDK3 (K_i_ 8mM) (Bowker-Kinley *et al*, 1998). The diversity of results from DCA treatment may relate to the expression profile of the PDKs.

The present study is the first to consider the contribution of all four PDKs towards the Warburg effect and the targeting of the PDK-PDH axis using DCA.

## Materials and Methods

### Cell culture

Sources of the cell lines and special media are detailed in the supplementary materials. All cells were grown at 37°C, 5% CO_2_ in RPMI media supplemented with 10% fetal bovine serum (FBS), 10 mM HEPES and 2 g/L NaHCO_3_ with the exception of MCF10A and V14 cells (see Suppl Methods for details). The V14 cancer cell line was derived by Dr. Anneke Blackburn from a spontaneous mammary adenocarcinoma arising in a BALB/c *Trp53^+/-^*mouse (Suppl Methods and (Blackburn *et al*, 2004)). For hypoxic conditions, cells were grown in a sealed humidified gas chamber (Billups-Rothenberg, Del Mar, CA, USA). Oxygen was flushed out with 95% nitrogen, 5% CO_2_ at 20 L/min for 4 min and gas inside flasks/plates was allowed to equilibrate for 3 hr. The chamber was then re-flushed as necessary to obtain O_2_ levels of 0.2-0.3%.

### *In vivo* tumor treatment with DCA

Animal experiments were conducted with the approval of the Australian National University Animal Ethics Experimentation Committee under the guidelines established by the Australian National Health and Medical Research Committee. Female BALB/c mice, 8-14 weeks old, were injected subcutaneously with 5 x 10^6^ V14 cells or 1 x 10^5^ 4T1 cells, and tumor size monitored with calipers. Established V14 tumors were treated with DCA, whereas 4T1-recipient mice were pretreated for 3 days with DCA, which did not affect the early development of these tumors (see Suppl methods for further explanation). DCA in the drinking water (1.5 g/L) delivered approximately 115 or 160 mg/kg/day for the 4T1 and V14 experiments, respectively. Tumor tissues were fixed overnight in neutral buffered formalin, and processed for H&E staining. The number of mitotic figures was counted per 10 high powered fields (hpf) per tumor.

### *In vitro* viability, apoptosis and proliferation assays

Neutral red staining was used to determine total viable cell number after 1-6 days of treatment, as described previously (Sun *et al*, 2011). Apoptosis was quantified by flow cytometry using FITC-labeled Annexin V and propidium iodide staining as previously described (Sun *et al*, 2011). Note that cells floating after drug treatment were collected and included in the analysis. Caspase 3/7 activities in were assessed by using Caspase-Glo 3/7 assay (Promega Corp., Madison, WI) according to the manufacturer’s instructions. Proliferation was quantitated using 5 µM carboxyfluorescein succinimidyl ester (CFSE) as previously described (Sun *et al*, 2011).

### Immunoblotting

Cells at ∼70% confluence were incubated in either normoxia or hypoxia for 18 hr. Cells were lysed with RIPA buffer (150 mM NaCl, 1% Triton X-100, 0.5% sodium deoxycholate, 0.1% SDS, 50 mM Tris, pH 8.0), sonicated (low power for 5 s) and centrifuged (13000 x g, 15 min, 4°C). Supernatant protein content was measured using BCA Assay Kit (Pierce, Rockford, IL, USA). Proteins (30 µg/sample) were separated via reducing 10% SDS-PAGE and standard western blotting procedures (Lim *et al*, 2004) were used to detect proteins of interest with the following primary antibodies: PDK1 (Cat# ADI-KAP-PK112, Enzo Life Sciences, Plymouth Meeting, PA, USA), PDK2 (Cat# AP7039b, Abgent, San Diego, CA, USA), PDK3 (Cat# H00005165-M01, Abnova, Taiwan), PDK4 (Cat# AP7041b, Abgent), β-actin (Abcam), HIF1α (Cat# NB100-134, Novus, Cambridge, UK).

### Extracellular lactate

Cells were seeded into 12-well plates (2×10^5^ for mouse lines, 1×10^5^ for human lines). After 24 hr, media was removed and DCA was added with fresh media. After 4 hr (mouse lines) or 24 hr (human lines), media was collected for lactate measurement as previously described (Sun *et al*, 2010). Lactate production at 24 hr was standardized to protein content to account for changes in cell number.

### PDH phosphorylation

Levels of pSer232, pSer293, and pSer300 were measured in cells treated with DCA for 3 hrs using the phosphoPDH In-Cell ELISA kit according to the manufacturers protocol (#MSP48, MitoSciences).

### PDK gene silencing with RNAi

Cells were transfected with 10 nM PDK siRNA (siPDK) or negative control siRNA (siNC) using Lipofectamine RNAiMAX (Invitrogen Lifesciences) following the manufacturers protocol. After 48 hr, knockdown efficiency was determined by immunoblotting (≥70% knockdown achieved), and drug treatment commenced. The total number of viable cells was measured 48 hr later.

### Gene expression database analysis using Oncomine

Data available through the Oncomine microarray gene expression analysis tool (www.oncomine.org) was analysed for expression of PDK1-4 in various cancer types. Differential analysis of mRNA expression levels in cancer compared to normal tissue in the same dataset was performed by the Oncomine resource, and the fold change and p values (Student’s t test with correction for multiple hypothesis testing) reported (Rhodes *et al*, 2007). To determine the proportion of cancers that overexpressed the PDK, the Oncomine waterfall plots were examined and the number of cases with expression greater than 2-fold the median expression level in normal tissues were counted (i.e. 1+log 2 (median centered-ratio)). Datasets examined were: TCGA Breast - including normal breast (n=61), invasive breast carcinoma (IBC, n= 76), invasive ductal breast carcinoma (IDBC, n=392), invasive lobular breast carcinoma (ILBC, n=36); TCGA Colorectal - including normal colon (n=19), colon adenocarcinoma (ColCa, n=101); Taylor Prostate 3 (taylor 2010) – including prostate gland (n=29), prostate carcinoma (ProstCa, n=131); Pei Pancreas (peiH09) – including normal pancreas (n=16), pancreatic carcinoma (PanCa, n=36), TGCA Brain – including brain (n=10), brain glioblastoma (glioblast, n=515).

### Statistical analysis

Differences between groups were analyzed with one-way ANOVA, comparing all treatment groups with each other using the Bonferonni post-hoc test in GraphPad Prism software. Each independent experiment was performed with at least three triplicate samples per treatment group. All results are expressed as mean±SEM of replicate values from at least two independent experiments. Minimum significance was at p<0.05.

## Results

### DCA can inhibit tumor growth *in vivo*

To examine the effectiveness of DCA *in vivo* against primary mammary tumors, two mouse mammary tumor cell lines were investigated both *in vitro* and *in vivo*. After 3 days of oral treatment with DCA, growth of established V14 tumors was halted (Figure 1A). This growth-arrested state was maintained for 2 weeks of treatment, with no regression. Histologically, the number of mitotic figures was significantly reduced by 35% (p=0.04) but there was no sign of apoptosis (Figure 1C-E). DCA treated V14 tumors also tended to display a greater degree of inflammation than control tumors (Supplementary Figure 1). In contrast, 4T1 tumors showed little response to DCA *in vivo* (Figure 1B and E).

**Figure 1.**
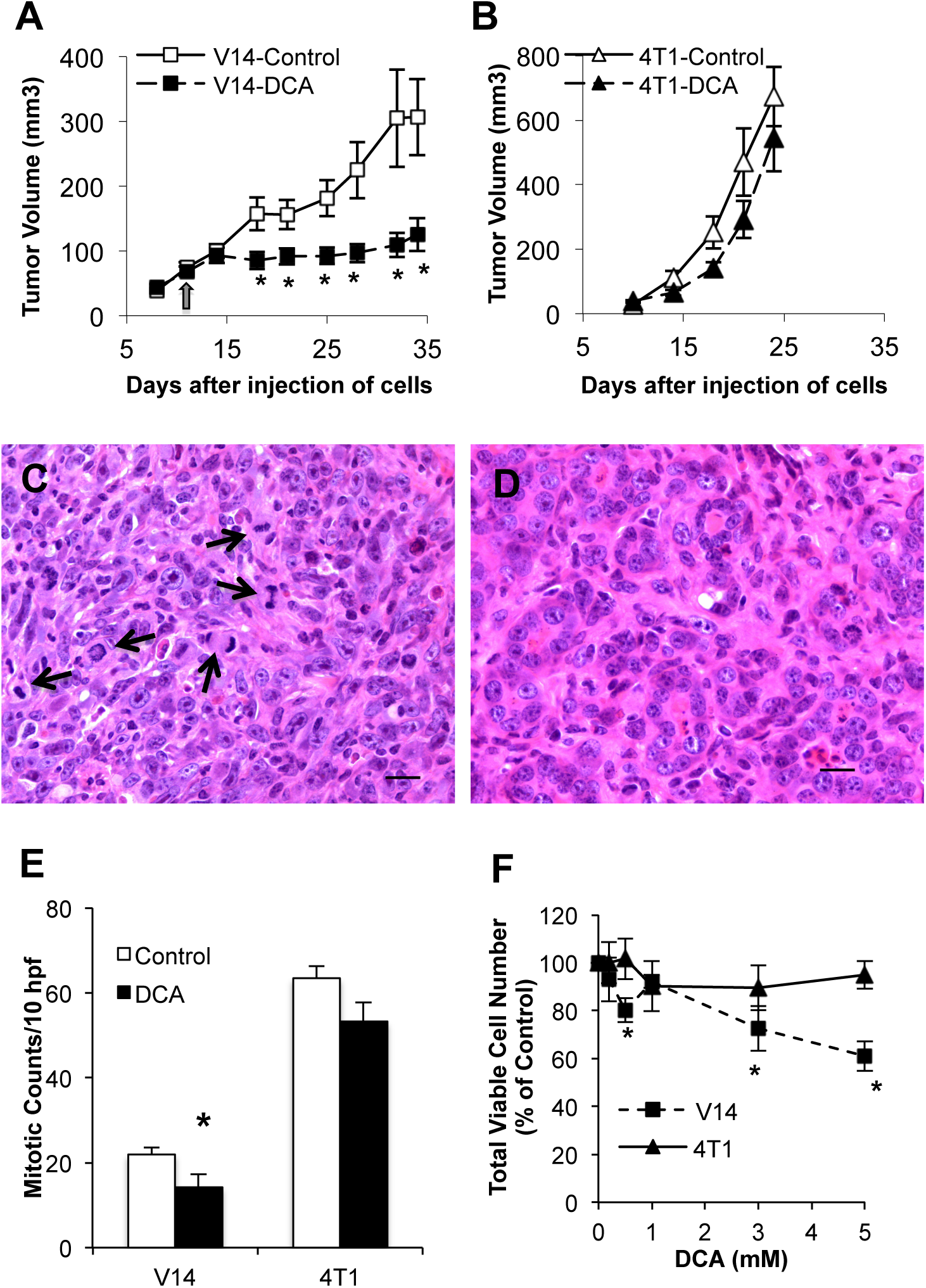
Inhibition of tumor growth *in vivo* by DCA. (A) Inhibition of subcutaneous V14 tumor growth *in vivo*. Arrow indicates the start of DCA treatment of established tumors. *p<0.05 vs untreated tumors (n=12 and 10 for control and DCA treated groups respectively). (B) DCA treatment did not inhibit 4T1 tumor growth *in vivo* (n=5 per group). Histology of V14 tumors untreated (C) or DCA treated (D) demonstrating decreased mitotic figures (arrows) with DCA treatment (photomicrographs x630, scale bar=20um), quantified in (E). (F) Total viable cell number after 48 hr DCA treatment *in vitro* in 4T1 and V14 cells. *p<0.05 and **p<0.005 vs untreated cells.

### DCA inhibits cancer cell growth without inducing apoptosis

The V14, 4T1 and a panel of human cancer cell lines were tested in vitro to investigate further the effects of targeting the PDK-PDH axis with DCA. *In vitro*, the malignant 4T1 cell line showed very little growth inhibition by DCA, whereas the non-metastatic V14 tumor line was highly sensitive to DCA, with 39% reduction in total cell number being achieved with 5 mM for 48 hr (Figure 1F). This effect on V14 cells was even more pronounced when treatment was maintained for up to 6 days, yet 4T1 growth was still uninhibited (Supplementary Figure 2). Treatment of a range of human cancer cell lines representing four major cancer types (breast, colon, pancreatic and prostate), found that DCA treatment over 48 hr reduced total viable cell number in 6/7 cancer cell lines, but not the non-cancerous MCF10A cells (Figure 2A). However, the extent of growth inhibition was variable, ranging from 5-30%. T-47D cells were the most sensitive. The remaining cancer cell lines were grouped as having moderate or low sensitivity to DCA treatment, based on their response across several DCA concentrations.

**Figure 2.**
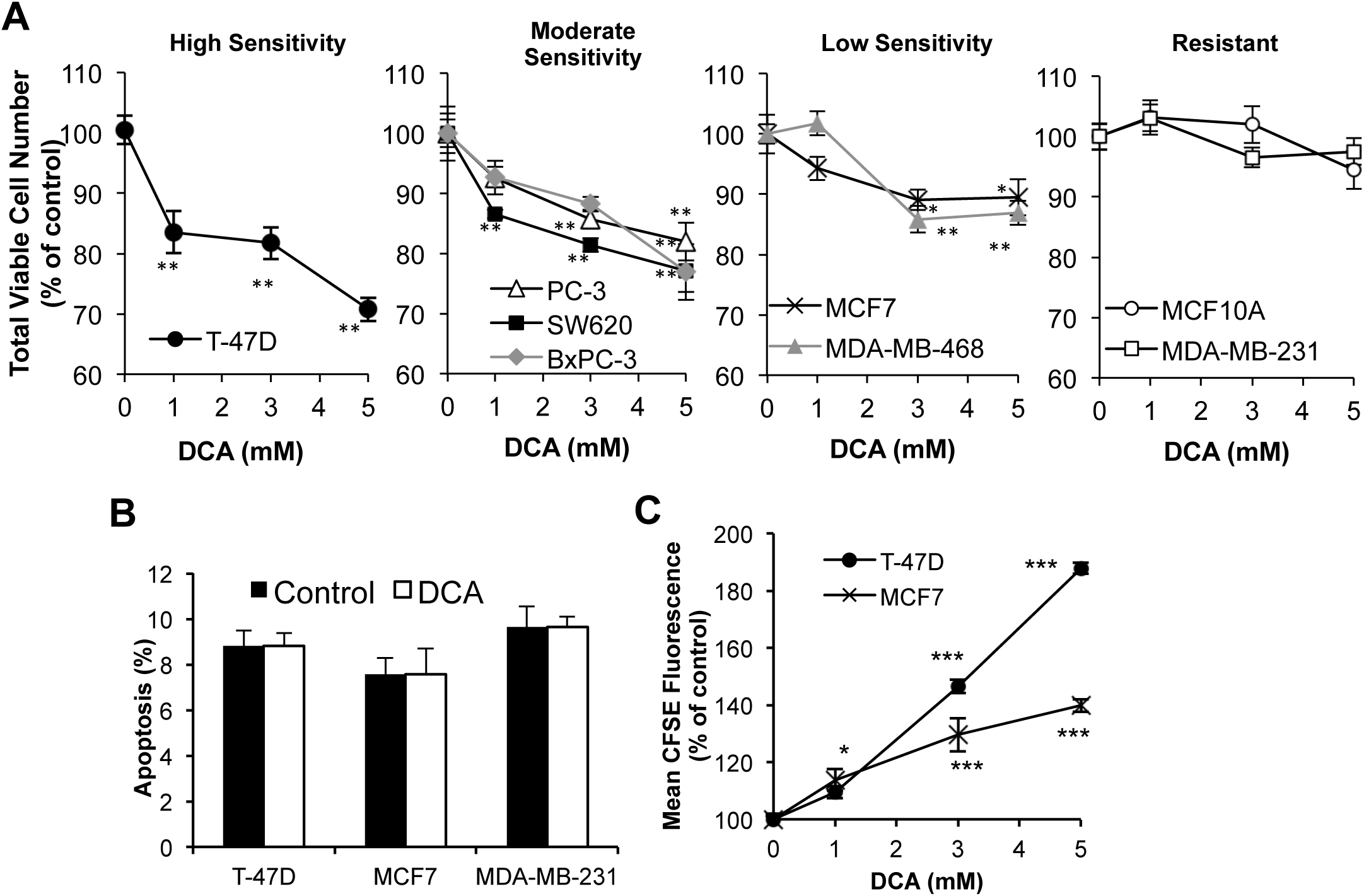
DCA inhibited proliferation. (A) Total viable cell number after 48 hr DCA treatment in human breast (T-47D, MCF7, MDA-MB-231, MDA-MB-468), colon (SW620), prostate (PC-3) and pancreatic (Bx-PC3) cancer cells, with MCF10A as a non-cancerous control. Cell lines are grouped into high, moderate or low sensitivity and resistant. (Note: y-axes start at 60%.) (B) Annexin V positive (apoptotic) cells after 48 hr of 5 mM DCA treatment. 1 µM staurosporine induced >60% apoptosis in all three cell lines (results not shown). (C) Mean CFSE fluorescence in DCA treated T-47D and MCF7 cells after 72 hr. *p<0.05, **p<0.01 and ***p<0.001 vs untreated control.

Morphologically, no apoptosis was apparent. Measurement of caspase 3/7 activity in V14 and T-47D cells could detect no significant increase after DCA treatment (Supplementary Figure 3), similar to our previous finding in 13762 MAT rat mammary adenocarcinoma cells (Sun *et al*, 2010). Three human breast cancer cell lines of varying sensitivity to DCA (T-47D, MCF7 and MDA-MB-231) were further tested for apoptosis by annexin V staining and FACS analysis. DCA did not induce apoptosis (Figure 2B) indicating DCA primarily inhibits proliferation, consistent with the effect seen *in vivo* (Figure 1; (Sun *et al*, 2010)). Growth inhibition by DCA was confirmed using CFSE staining where greater fluorescence intensity, indicating reduced proliferation, was seen with DCA treatment in T-47D and MCF7 cells (Figure 2C). The ∼2-fold increase in CFSE fluorescence after 72 hrs in T-47D cells (indicating one less population doubling than untreated cells) is consistent with the ∼30% lower cell number at 48 hrs (which could reasonably extrapolate to 50% at 72 hrs), further suggesting that inhibition of proliferation could account for the major part of the effects of DCA in these experiments.

### Growth inhibition by DCA is determined by the PDK expression profile

DCA targets the family of four PDKs, which are inhibited by concentrations of DCA ranging from 0.2 – 8.0 mM (Bowker-Kinley *et al*, 1998). Thus the expression of all four PDK isoforms in these cancer cells was investigated to determine if the PDK profile correlated with sensitivity to DCA. An ideal PDK profile for DCA inhibition would be expression of PDK2 (lowest K_i_ for DCA at 0.2 mM) and low levels of the other isoforms, in particular PDK3 (highest K_i_ of 8 mM). Immunoblotting in the panel of cancer cell lines revealed that PDK2 was expressed in all seven cell lines, however expression of the other three isoforms varied. As expected, T-47D cells expressed predominantly PDK2 with low levels of the other isoforms in normoxia (Figure 3A), correlating with them being most sensitive to growth inhibition by DCA.

**Figure 3.**
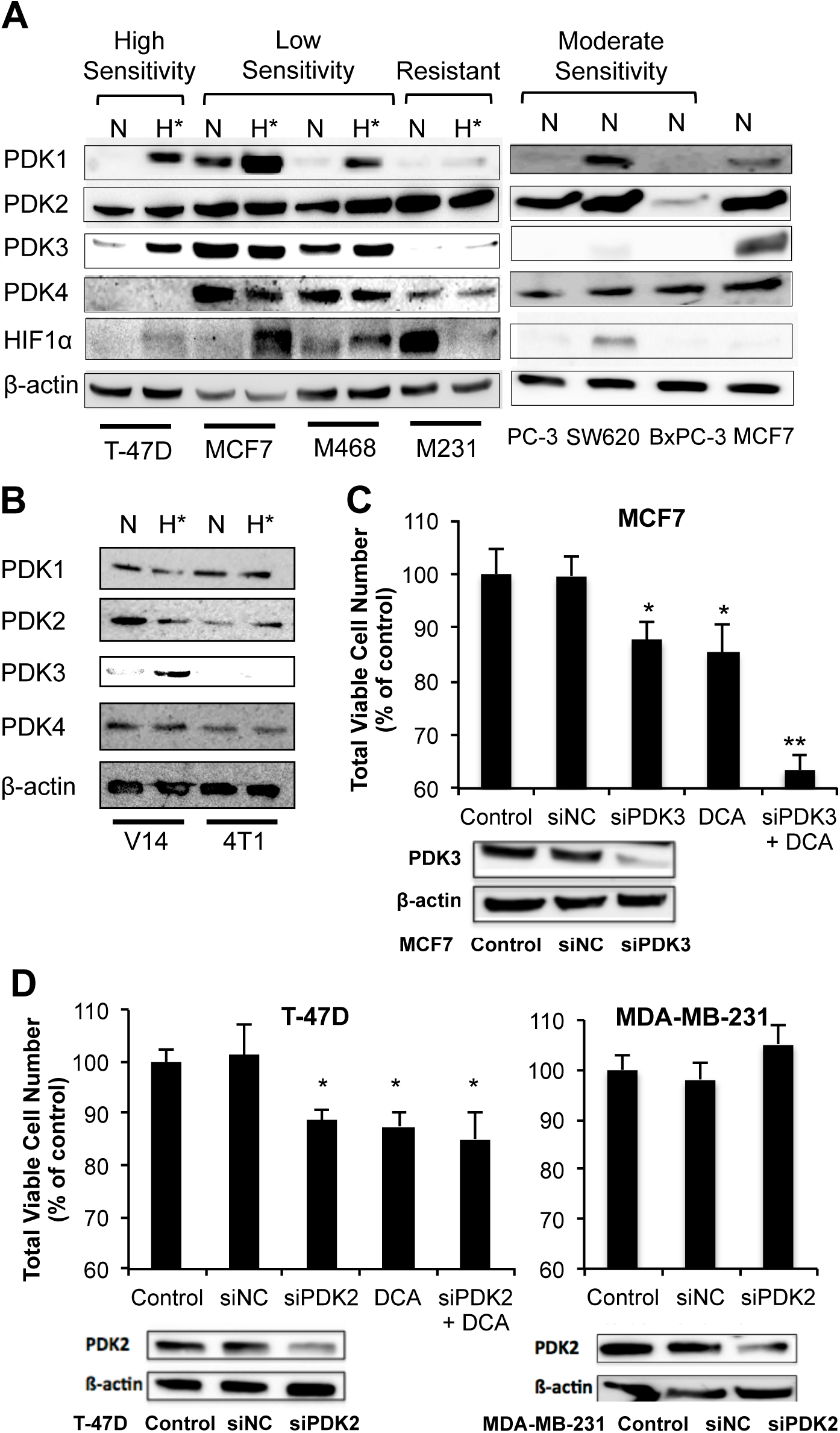
Expression of PDK isoforms varies across cancer cell lines and determines sensitivity to DCA growth inhibition. (A) Expression of PDK1-4 and HIF1α in human breast, colon (SW620), prostate (PC-3) and pancreatic (Bx-PC3) cancer cells. Cell lines are grouped into high, low or moderate sensitivity or resistant to growth inhibition by DCA, with MCF7 cells as a comparison point across different blots. N, normoxia; H*, hypoxia (0.2% O_2_). (M468 – MDA-MB-468, M231 – MDA-MB-231). (B) Expression of PDK1-4 in mouse mammary tumor cell lines in normoxia (N) and hypoxia (H*). (C) Total viable cell number after 48 hr of 5 mM DCA treatment in PDK3 knockdown (siPDK3) MCF7 cells. siNC – negative control siRNA. Blot shows PDK3 protein expression 48 hr post-transfection with siRNA. (D) Total viable cell number 48 hr after knockdown of PDK2 by siPDK2 transfection of T-47D (+/− 1 mM DCA) and MDA-MB-231 cells. Blot shows PDK2 protein expression 48 hr post-transfection with siRNA. *p<0.05 and **p<0.01 vs untreated control. (Note: y-axes start at 60%.)

The relative sensitivity to DCA of 5 out of the other 6 human cell lines investigated also correlated with the PDK expression profile. In normoxia, the moderately sensitive BxPC-3, SW620 and PC-3 cells expressed PDK2, however they also expressed additional isoforms: PDK4 in all 3 cell lines and PDK1 in SW620 cells (Figure 3A). MCF7 and MDA-MB-468 cells, which displayed low sensitivity to DCA, expressed high levels of PDK3 in normoxia, with MCF7 cells also expressing high levels of PDK1. MDA-MB-231 cells were anomalous, as their profile was like T-47D cells, but they displayed low sensitivity to DCA. The profile of mouse V14 cells (Figure 3B) showed low levels of PDK3 expression in normoxia (but which could be induced in hypoxia), consistent with moderate to high sensitivity to DCA, whereas 4T1 cells were anomalous like MDA-MB-231 cells, lacking overexpression of PDKs yet showing low / insensitivity to DCA. Of note, both MDA-MB-231 and 4T1 cells are triple negative breast cancer cell lines.

To confirm sensitivity to DCA growth inhibition was determined by the PDK expression profile, the highest K_i_ isoform, PDK3, was knocked down using siRNA in MCF7 cells. This resulted in an increase in the effect of 5 mM DCA from 13% to 37% decrease in viable cell number, comparable to the high sensitivity line T-47D (Figure 3C). Further, targeting PDK2 by either inhibition with 1 mM DCA or knockdown with siPDK2 (∼70%) led to similar effects on growth inhibition in T-47D cells (Figure 3D). The addition of 1 mM DCA on siPDK2 T-47D cells resulted in no further decrease in growth, confirming that DCA inhibited growth through PDK2 inhibition. Thus the ability of DCA to inhibit cancer cell growth was due to on-target PDK inhibition. As MDA-MB-231 cells were resistant to growth inhibition by DCA (Figure 2A), it was expected that there would be little reduction in viable cell number from decreased PDK2 activity. This was indeed the case, with no reduction in viable cells occurring after siRNA knockdown of PDK2 (Figure 3D).

### DCA growth inhibition coincides with activation of PDH and reversal of the Warburg effect

To further confirm DCA was acting on-target, and to address the difference between cell lines that were sensitive (T-47D and MCF7) or resistant (MDA-MB-231) to growth inhibition by DCA, the biochemical responses to DCA were investigated. As DCA inhibits the PDKs, it should reduce phosphorylation of PDH at Ser232, 293 and 300 (indicators of PDH activity) and reduce levels of extracellular lactate (indicator of the Warburg effect). This was the case in T-47D and MCF7 cells, indicating PDH activation and reversal of the Warburg effect, but not in the resistant MDA-MB-231 cells (Figure 4A and 4B). T-47D cells showed the largest decrease in phosphorylation at all 3 sites after 1 mM DCA, consistent with their PDK profile and high sensitivity to DCA growth inhibition, with MCF7 cells also showing a significant decrease in phosphorylation in 2/3 sites at a higher (5 mM) concentration of DCA (Figure 4A).

**Figure 4.**
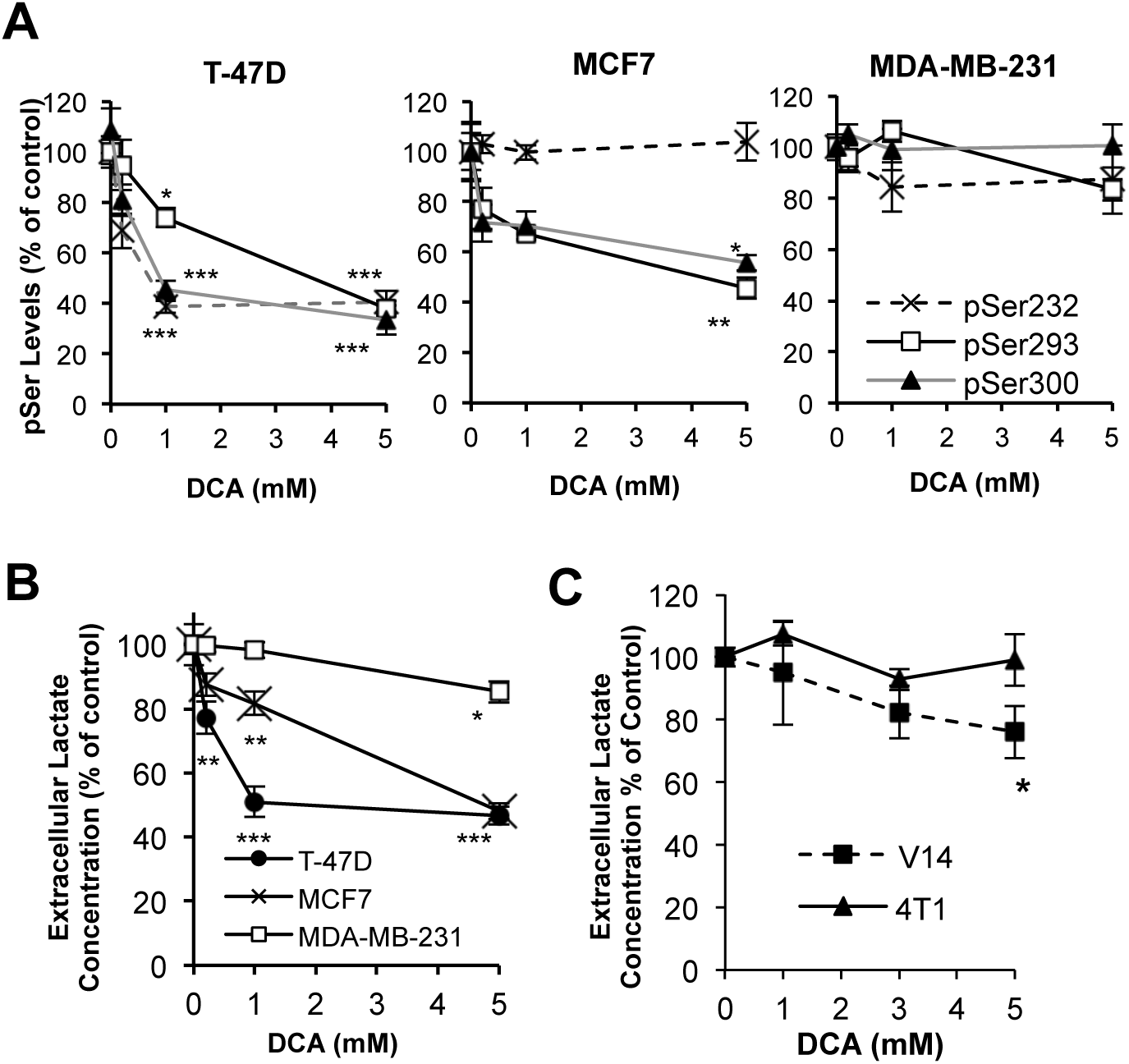
DCA activated PDH, reversed the Warburg effect and inhibited growth via PDK inhibition. (A) Levels of pSer232, pSer293, pSer300 after 3 hr DCA treatment in T-47D, MCF7 and MDA-MB-231 cells. (B) Extracellular lactate levels in human breast cancer cell lines after 24 hr DCA treatment. (C) Reduction of 4 hr extracellular lactate production by DCA in mouse mammary tumor cell lines. *p<0.05 compared to corresponding 4T1 cells. *p<0.05, **p<0.01, ***p<0.001 vs untreated control.

Consistent with the PDH phosphorylation, measurements of lactate production found 24 hr DCA treatment at 1 mM was sufficient to achieve maximal decrease in lactate production in T-47D cells (49% decrease), with a similar effect also being achieved in MCF7 cells at 5 mM (Figure 4B). Lactate production was also significantly reduced in the sensitive V14 cells indicating reversal of the Warburg effect, but not in the resistant 4T1 cells (Figure 4C). In MDA-MB-231 cells, there was only ∼10% reduction in lactate and pSer levels after DCA treatment (Figure 4A & 4B) in keeping with the lack of growth inhibition (Figure 2A). Thus in MDA-MB-231 and 4T1 cells, other enzymes are potentially dominating the fate of pyruvate metabolism over that of PDH.

### DCA can induce apoptosis in hypoxia

PDK1 and 3 expression are up-regulated in hypoxia (Lu *et al*, 2008; Papandreou *et al*, 2006) and this may reduce the effectiveness of DCA *in vivo*. Conversely, we and others have shown that DCA can reduce HIF1α expression (Sun *et al*, 2011; Sutendra *et al*, 2012; Supplementary Figure 4) which may decrease cell viability in hypoxia. Thus we investigated the effects of hypoxia on the PDK expression profile and on the growth and viability of breast cancer cells in the presence and absence of DCA. Exposure to hypoxia (∼0.2% oxygen) for 48 hr robustly induced the expression of PDK1 and PDK3 in most cells where expression was low in normoxia, and this was accompanied by HIF1α stabilization (Figure 3A). Where PDK1 or PDK3 was expressed in normoxia, a further increase in PDK was also detectable in hypoxia for PDK1 in MCF7 cells and PDK3 in MDA-MB-468 cells, however there was no further increase in PDK3 expression in MCF7 cells. MDA-MB-231 cells did not induce HIF1α or PDK protein expression during hypoxia. In fact, HIF1α levels decreased in hypoxia and despite the high levels of HIF1α in normoxia, MDA-MB-231 cells did not express PDK1 or PDK3, indicating a very dysfunctional HIF1/PDK pathway.

Hypoxia reduced the viable cell number in all cell lines after 48 hr and DCA had no additional effect on cell number in 3 of the cell lines (Figure 5A). This is consistent with DCA acting via the inhibition of cell growth, and thus being unable to further reduce cell number in the growth-inhibited hypoxic environment. However DCA was able to further reduce viable cell number in MCF7 cells in hypoxia compared to hypoxia alone (Figure 5A) with a 3-fold increase in apoptotic MCF7 cells compared to hypoxia alone (Figure 5B).

**Figure 5.**
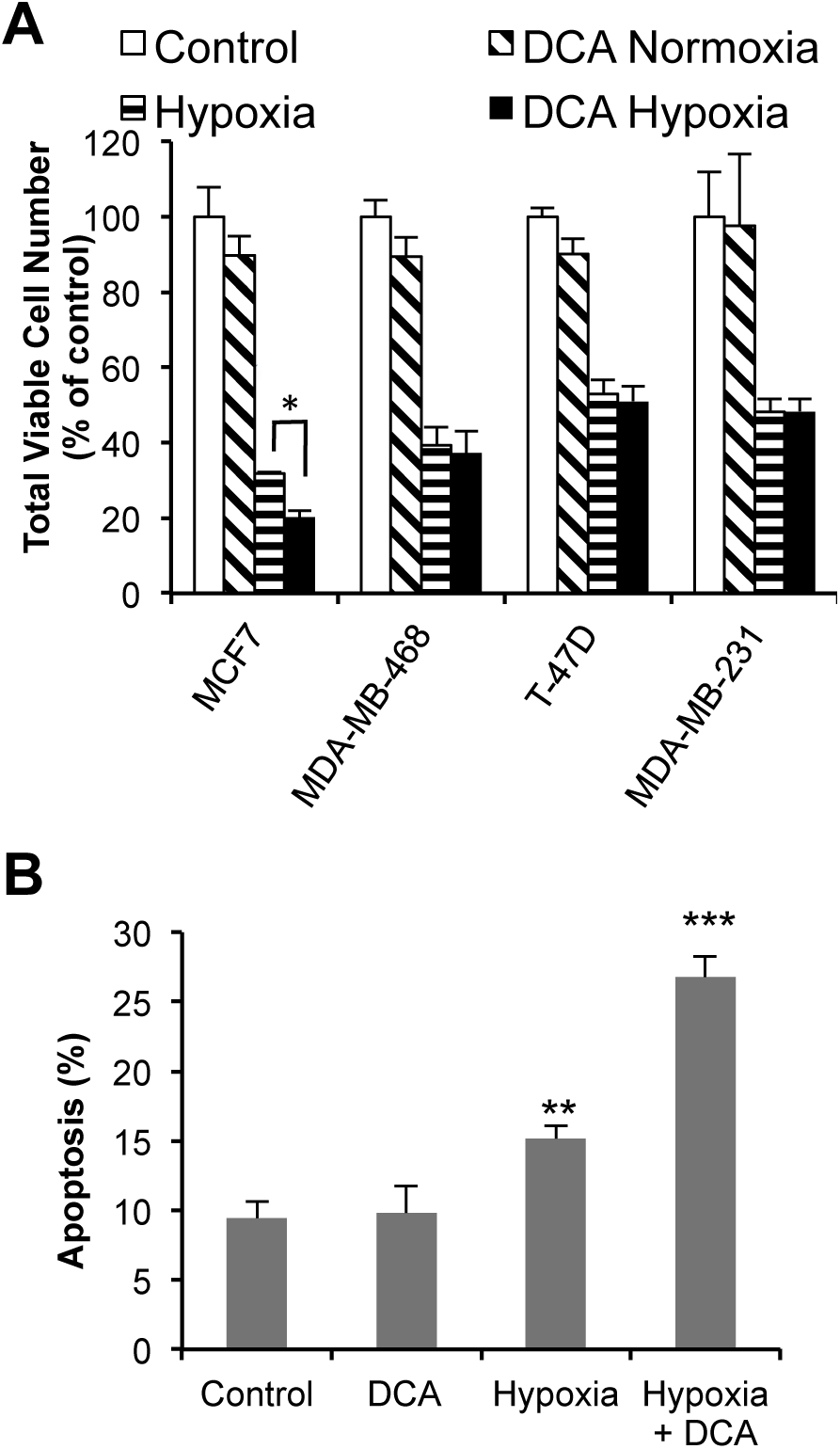
DCA induced apoptosis in hypoxia. (A) Growth of breast cancer cells in normoxia and 0.2% O_2_ over 48 hr +/− 3 mM DCA. (B) Annexin V positive (apoptotic) MCF7 cells after 48 hr growth +/− 3 mM DCA in normoxia or hypoxia. *p<0.05, **p<0.01 vs control, ***p<0.001 vs control.

### PDK mRNA Profiles in Human Cancers

To examine whether PDK expression also varied in patient tumors, the PDK RNA expression profiles of common cancer types were examined using the Oncomine database tool (Rhodes *et al*, 2007). Comparing cancer expression levels to the corresponding normal tissue in the same dataset (Figure 6 and Supplementary Table 1), PDK3 mRNA was significantly overexpressed in breast cancer (2.3-fold, p=1.19 x 10^-24^) and colon cancer (1.5-fold, p=9.24 x 10^-8^), with 45% of breast and 20% of colon cancers showing greater than 2-fold the median normal mRNA expression level (Supplementary Table 2). Thus a considerable fraction of breast and colon cancers may be unresponsive to DCA treatment due to PDK3 overexpression. In contrast, glioblastoma, prostate and pancreatic cancers showed lower mRNA expression of PDK3 than their normal tissue counterparts, suggesting that these cancer types may be good candidates for treatment with DCA. PDK1 was not markedly overexpressed in the cancers examined (Figure 6). However, on closer examination it was noted that 23% of IDBC samples and 15% of glioblastoma samples overexpressed PDK1 mRNA (Figure 6 and Supplementary Table 2), thus PDK1 expression may also contribute to low sensitivity to DCA treatment in a proportion of these cancers.

**Figure 6.**
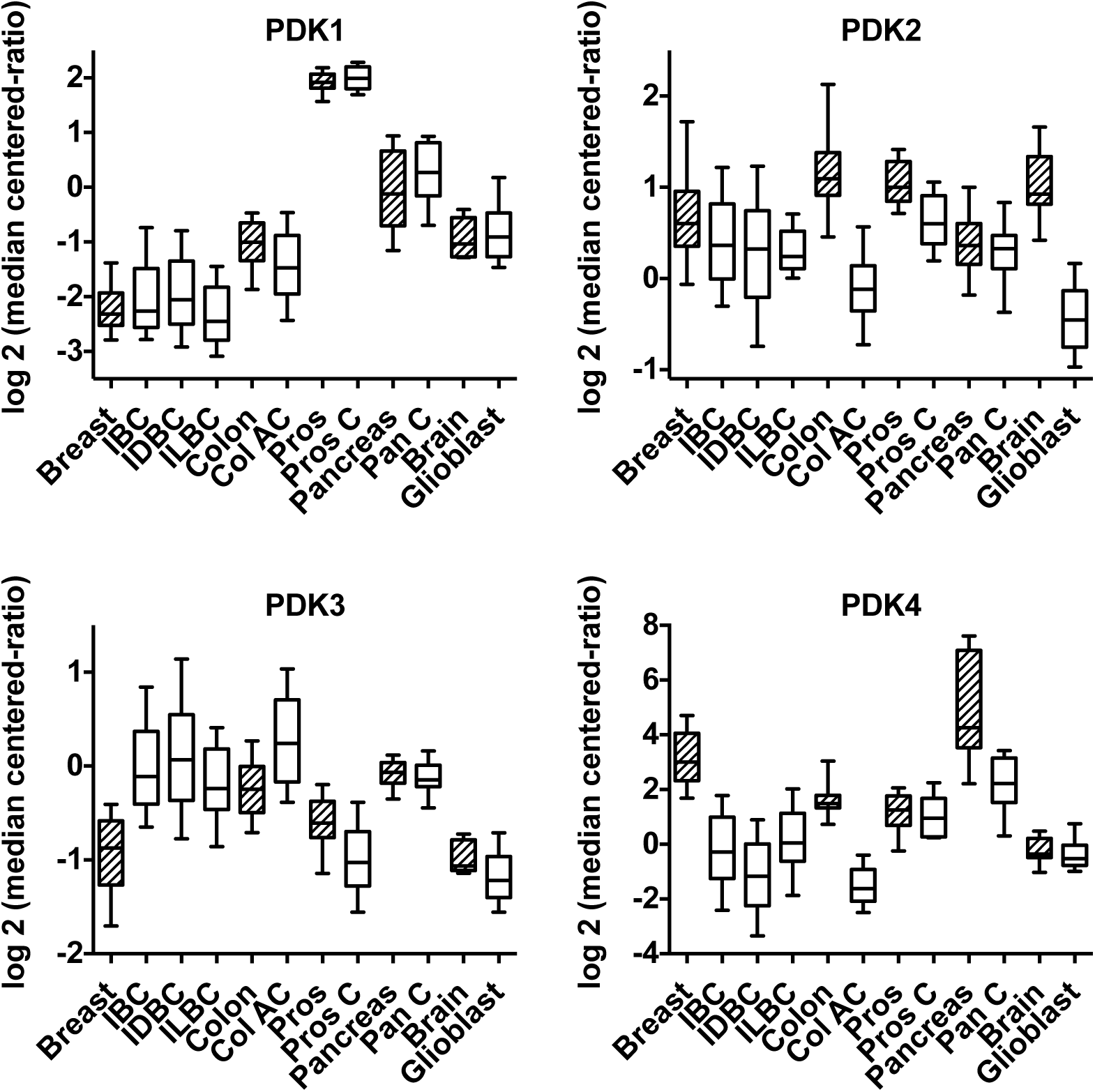
Expression of PDK1-4 mRNA in human cancer tissues. Datasets available through Oncomine were examined for PDK expression. Graphs display median (line), 25-75 percentiles (box), and 10-90 percentiles (whiskers). Shaded boxes represent normal tissues with the corresponding cancer tissue following in open boxes. Abbreviations in methods. Statistical data in Supplementary Table 1.

## Discussion

The generality of the Warburg effect across many cancers offers hope that manipulating tumor metabolism will be clinically useful in cancer treatment. DCA is a compound that can reverse the Warburg effect and has clinical potential due to its previous history of use in congenital lactic acidosis (Stacpoole & Greene, 1992; Stacpoole *et al*, 1983), and is currently undergoing phase I/II clinical trials in several centers against a range of cancer types (clinicaltrials.gov). In our study, DCA was effective against a mouse primary mammary tumor model, halting growth of established tumors *in vivo* (Figure 1). Further DCA was effective in inhibiting growth of 6/7 human tumor cell lines from common cancers (Figure 2). The relative effectiveness of DCA correlated with its on-target actions (reversing the Warburg effect and reducing PDH phosphorylation, Figure 1B and 4) and with the expression profile for the four PDK isoforms (Figure 3), which require increasing concentrations of DCA for inhibition (PDK2 < PDK4 < PDK1 << PDK3). Collectively the findings indicate that all four PDK isoforms should be considered when designing PDK inhibitors.

To determine whether PDK expression profiles also varied in human tumors, and thus whether the PDKs are potential biomarkers in the clinic, the Oncomine database was consulted for PDK mRNA expression of patient tumors and compared with published immunohistochemical studies. Immunohistochemistry of 74 paired normal and colorectal cancer tissues reported up-regulation of PDK3 in cancer of all stages that was associated with earlier relapse, while PDK1 levels decreased (Lu *et al*, 2011). This discordant regulation of PDK1 and PDK3 is in agreement with our Oncomine findings for colon cancer (Figure 6). Michelakis et al report on PDK protein levels in a small number of glioblastoma tissues (n=8) and patient-derived primary cell lines (n=3) by western blotting. For PDK1, protein and Oncomine results correlated well, with 2/8 tissues showing high PDK1 protein vs 15% of TCGA glioblastoma samples having high mRNA levels (Supplementary Table 2). The complete lack of overexpression of PDK3 in the TCGA and several other Oncomine glioblastoma datasets also correlates with the apparent positive outcomes for treating glioblastomas *in vitro* or *in vivo* with DCA (Michelakis *et al*, 2010). Thus it appears the mRNA expression levels of PDKs obtained from Oncomine (Figure 6) are indicative of PDK protein levels in human tumors. Given the varying PDK expression in cancers from the Oncomine data (Figure 6), the PDK expression profiles are likely to vary greatly in human cancers as we observed in cell lines, thus the PDK expression profiles are potential biomarkers for DCA sensitivity.

Of note, the two cancer cell lines that were resistant to growth inhibition by DCA (4T1 and MDA-MB-231; Figure 2) are both triple-negative (ER-, PR-, HER2-) breast cancer cell lines. Biochemical analysis revealed that in MDA-MB-231 cells, DCA was unable to activate PDH to redirect pyruvate metabolism away from lactate and towards acetyl-CoA production (Figure 4). This indicates that other enzymes involved in pyruvate metabolism may be over-expressed in triple-negative breast cancer cells, driving the production of lactate. This is indeed the case, as it has recently been shown that in a panel of 23 triple-negative breast cancer cell lines, 20 expressed lactate dehydrogenase B (LDHB) as well as the commonly expressed LDHA (McCleland *et al*, 2012). This suggests that there is greater overall LDH activity in triple-negative cells, which may be indicative of a metabolic phenotype that, through multiple changes in gene expression, renders cells resistant to changes in flux through PDH and therefore insensitive to DCA. The MDA-MB-468 are also triple-negative but were sensitive to DCA growth inhibition (Figure 2), however they were one of the three triple-negative cell lines that did not express LDHB (McCleland *et al*, 2012). Thus the PDK expression profile may not be a suitable biomarker for DCA sensitivity in triple-negative breast cancers.

As all four PDKs are involved in determining sensitivity to DCA, expression levels of PDKs are potential clinical biomarkers for DCA/PDK inhibitor sensitivity. Patient tumors with high expression of PDK1/3 may be less suitable for treatment with DCA. DCA is most likely to be used in combination with other agents, as elimination rather than growth arrest of cancer cells is generally the aim. However, there may be a role for using DCA therapy alone to manage cancer growth in special circumstances, such as palliative care in the elderly (Khan, 2011). Thus harnessing the anti-proliferative potential of DCA as a single agent, particularly in cancers with low expression of PDK3, may be clinically useful.

Our study has focused on the anti-proliferative effects of DCA, as we did not observe increased apoptosis (Figure 2). While the initial study of the anti-cancer effects of DCA induced apoptosis as well as inhibiting proliferation (Bonnet *et al*, 2007), subsequent studies have often used unreasonably high concentrations of DCA to achieve a similar result, which is unlikely to be due to on-target effects. There are however many reports of reduced cell growth and proliferation with DCA treatment (Cao *et al*, 2008; Chen *et al*, 2009; Sanchez-Arago *et al*, 2010; Sun *et al*, 2010; Sun *et al*, 2009). This does not appear to involve a specific block in cell cycle, but rather a general increase in population doubling times (Cao *et al*, 2008; Stander *et al*, 2011; Washington & Quintyne, 2012) associated with tumor-specific changes to metabolic proteins (Stockwin *et al*, 2010). The present study demonstrates that when inhibiting proliferation, DCA is acting on target, modifying PDH phosphorylation at concentrations consistent with PDK profiles (Figure 4). The increased PDH activity would be expected to alter the flux through glycolysis, and hence the availability of intermediates upstream of PDH for pathways such as the pentose phosphate pathway (providing NADPH for fatty acid synthesis and redox control), and the serine synthesis pathway. Balancing these aspects of metabolism, as well as ATP availability, are critical for cancer cells to maintain their proliferation rate (Cairns *et al*, 2011). Thus we speculate that the anti-proliferative effects of DCA are due to a reduction in the availability of precursors necessary for biomass production and redox regulation during proliferation. Metabolomic analysis is currently underway to examine this hypothesis.

An alternative explanation for the range of responses to DCA is variable drug uptake. Babu et al reported that the apoptotic effects of DCA were limited by the uptake of DCA into cancer cells, which was overcome by increasing the expression of the plasma membrane transporter SLC5A8 (Babu *et al*, 2011). However, our ability to detect changes in PDH phosphorylation in 3 hr at low DCA concentrations is consistent with DCA rapidly accessing the cells and acting on PDKs to reduce lactate production (Figure 4A and B) without the need to induce expression of SLC5A8. We postulate that the increased apoptosis from SLC5A8 overexpression observed by Babu et al. is due not to improved PDK inhibition from increased uptake, but rather due to off-target effects of DCA, or to altered transport of other metabolites creating additional stress and synergizing with DCA.

Under the additional stress of hypoxia, we found that DCA induced apoptosis in MCF7 cells, but had no effect on total cell number in other cell lines (Figure 6). This difference in response may also be attributed to the PDK profiles. PDK1 and 3 are induced by hypoxia, however as MCF7 cells already displayed high levels of expression of these genes in normoxia, they were less able to adapt to the hypoxic stress than the other cell lines (Figure 3), and thus underwent apoptosis when challenged with DCA. Alternatively, decreased HIF1α levels may decrease survival during hypoxia (Sun *et al*, 2011; Sutendra *et al*, 2012;), however DCA was able to reduce HIF1α in both MCF7 and T-47D cells (Supplementary Figure 4), thus this would not seem to account for the different responses of these cell lines. *In vivo*, it may be have been expected that DCA would be less effective where areas of hypoxia would result in induction of PDK1 and 3, thus requiring higher concentrations of DCA to alter the PDH/PDK axis. However, our experience with V14 tumors found DCA was highly effective at inhibiting cell growth both *in vivo* and *in vitro* (Figure 1). This contrasts with a study on SW480 colon cancer cell lines (Shahrzad *et al*, 2010) where enhanced growth under hypoxia *in vitro* and *in vivo* were reported with DCA treatment. It will be important to determine the precise mechanism by which DCA alters the apoptotic responses of cancer cells, so that tumors which may undergo enhanced survival can be identified and excluded from DCA therapy.

PDKs are important regulators of metabolism in cancer, integrating signals from HIF1, MYC and growth factor receptors (Hitosugi *et al*, 2011; Kim *et al*, 2007; Lu *et al*, 2008; Papandreou *et al*, 2006). DCA is a prototype PDK inhibitor with clinical potential. Our study is the first to demonstrate that sensitivity towards PDK inhibitors is determined by the PDK expression profile of a tumor cell. The regulation of PDKs needs to be further investigated to allow for optimal outcome when treating patients with PDK inhibitors and for combining DCA/PDK inhibitors with additional chemotherapeutic agents.

## Acknowledgements

We thank Philip Board and Chris Parish for many helpful discussions. This project was co-funded through the PdCCRS (#1008861) by the National Breast Cancer Foundation and Cancer Australia, and by grants to ACB from Cancer Council ACT (#585409) and NHMRC Fellowship (#366787).

