## Supplemental Table 1 for "Pyruvate dehydrogenase kinase expression profile is a biomarker for cancer sensitivity to dichloroacetate-mediated growth inhibition"

Supplementary Table 1.

Differential expression analysis (normal vs cancer) of PDK1-4 in Oncomine datasets.

| | PDK1 | | PDK2 | | PDK3 | | PDK4 | |
|---|---|---|---|---|---|---|---|---|
| | p value | Fold change | p value | Fold change | p value | Fold change | p value | Fold change |
| **Overexpression** | | | | | | | | |
| Breast Cancer | | | | | | | | |
| IBC | 0.045 | 1.148 | 0.983 | -1.192 | **4.8E-16** | **1.971** | 1.000 | -5.718 |
| IDBC | **0.001** | **1.218** | 1.000 | -1.313 | **1.2E-24** | **2.292** | 1.000 | -9.088 |
| ILBC | 0.653 | -1.038 | 0.995 | -1.214 | **2.3E-10** | **1.799** | 1.000 | -5.002 |
| Colon Cancer | 0.566 | -1.015 | 1.000 | -1.650 | **9.2E-08** | **1.539** | 1.000 | -4.120 |
| Pancreatic Cancer | 0.044 | 1.316 | 0.704 | -1.057 | 0.080 | 1.259 | 1.000 | -7.074 |
| Prostate Cancer | 0.036 | 1.058 | 1.000 | -1.335 | 1.000 | -1.299 | 0.929 | -1.185 |
| Glioblastoma | 0.043 | 1.155 | 1.000 | -2.589 | **8.8E-06** | **1.331** | 0.555 | -1.017 |
| **Underexpression** | | | | | | | | |
| Breast Cancer | | | | | | | | |
| IBC | 0.012 | -1.223 | **1.6E-06** | **-1.423** | 1.000 | 1.550 | **2.8E-27** | **-10.905** |
| IDBC | **0.001** | **-1.221** | **5.2E-06** | **-1.325** | 1.000 | 2.168 | **1.6E-39** | **-20.098** |
| ILBC | **2.5E-05** | **-1.442** | 0.003 | -1.274 | 1.000 | 1.516 | **3.8E-14** | **-8.122** |
| Colon Cancer | 0.015 | -1.230 | **2.6E-12** | **-2.249** | 0.633 | 1.027 | **1.7E-18** | **-7.302** |
| Pancreatic Cancer | 0.921 | 1.214 | 0.296 | -1.057 | 0.145 | -1.050 | **2.0E-07** | **-5.194** |
| Prostate Cancer | 0.964 | 1.058 | **3.4E-08** | **-1.335** | **2.2E-06** | **-1.299** | 0.071 | -1.185 |
| Glioblastoma | 0.957 | 1.155 | **5.2E-07** | **-2.589** | **9.3E-04** | **-1.268** | 0.982 | 1.581 |
