## Supplemental Table 2 for "Pyruvate dehydrogenase kinase expression profile is a biomarker for cancer sensitivity to dichloroacetate-mediated growth inhibition"

Supplementary Table 2.

Proportion of cancer cases overexpressing (>2-fold normal) PDK1 or PDK3.

| | | PDK1 | | PDK3 | |
|---|---|---|---|---|---|
| | Total (n) | > 2-fold (n) | % over-expressing | > 2-fold (n) | % over-expressing |
| **TCGA Breast** | | | | | |
| Normal Breast | 61 | 4 | 6.6 | 1 | 1.6 |
| Breast Cancer | | | | | |
| IBC | 76 | 16 | **21.1** | 25 | **32.9** |
| IDBC | 392 | 91 | **23.2** | 178 | **45.4** |
| ILBC | 36 | 2 | 5.6 | 11 | **30.6** |
| **TCGA Colon** | | | | | |
| Colon | 19 | 0 | 0.0 | 0 | 0.0 |
| Colon Cancer | 101 | 5 | 5.0 | 20 | **19.8** |
| **Pei Pancreas** | | | | | |
| Pancreas | 16 | 2 | 12.5 | 0 | 0.0 |
| Pancreatic Cancer | 36 | 7 | 19.4 | 0 | 0.0 |
| **Taylor Prostate 3** | | | | | |
| Prostate Gland | 29 | 0 | 0.0 | 0 | 0.0 |
| Prostate Cancer | 131 | 0 | 0.0 | 1 | 0.8 |
| **TCGA Brain** | | | | | |
| Brain | 10 | 0 | 0.0 | 0 | 0.0 |
| Brain Glioblastoma | 515 | 76 | **14.8** | 8 | 1.6 |
