## Supplementary Figures for "Pyruvate dehydrogenase kinase expression profile is a biomarker for cancer sensitivity to dichloroacetate-mediated growth inhibition"

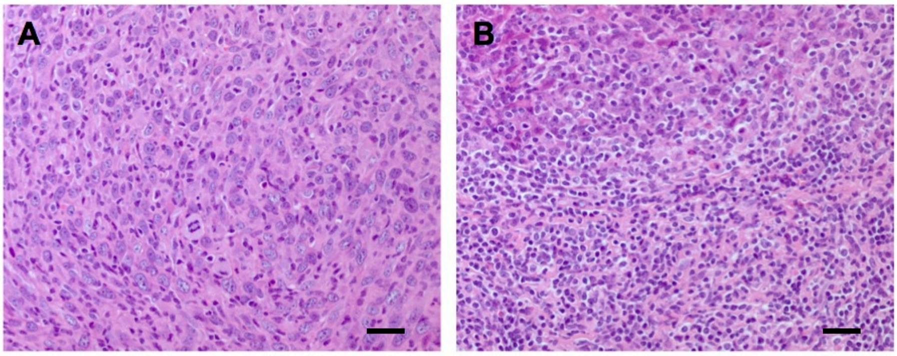


**Supplementary Figure 1**

Histology of V14 tumors untreated **(A)** or DCA treated **(B)** demonstrating increased lymphocyte infiltration (x400, scale bars=50um) with DCA treatment. 6/8 (75%) DCA treated vs 4/9 (44%) control tumors displayed moderate – severe inflammation, with lymphocytes more likely to be found throughout the tumor, and neutrophils more likely to be present within the tumor.

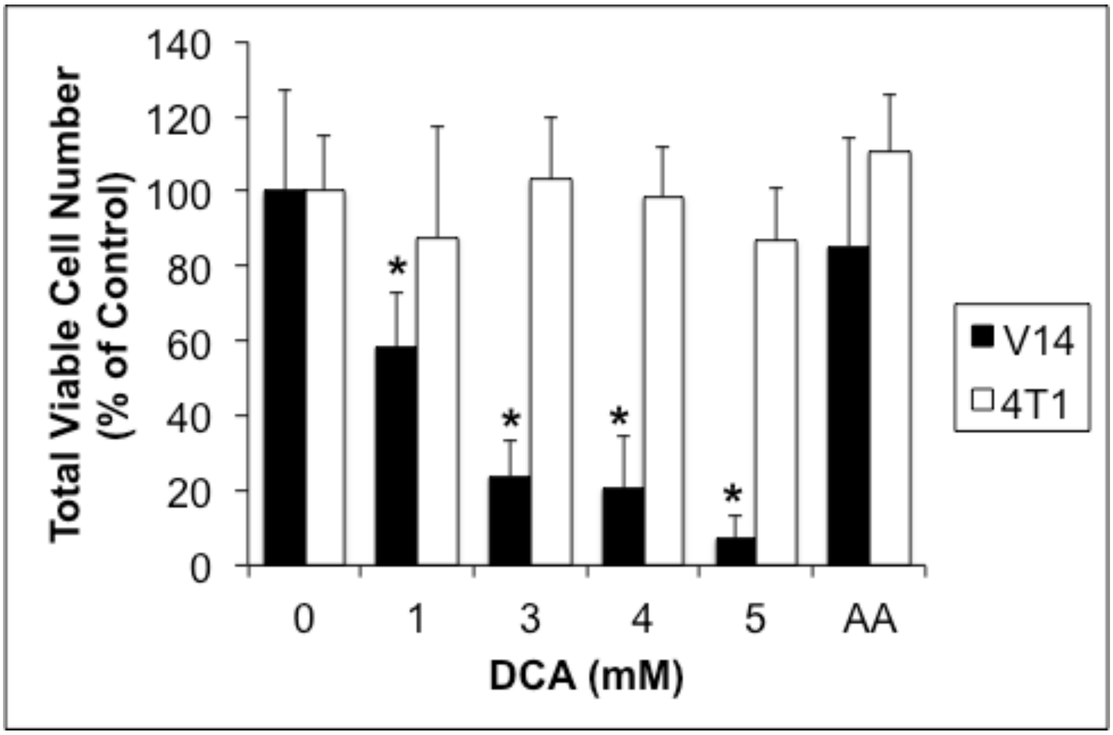


**Supplementary Figure 2**

Total viable cell number for mouse mammary tumor cell lines after 6 days of treatment with 0-5 mM DCA or 5 mM acetic acid (both neutralized). * p<0.005 compared to 0 mM DCA.

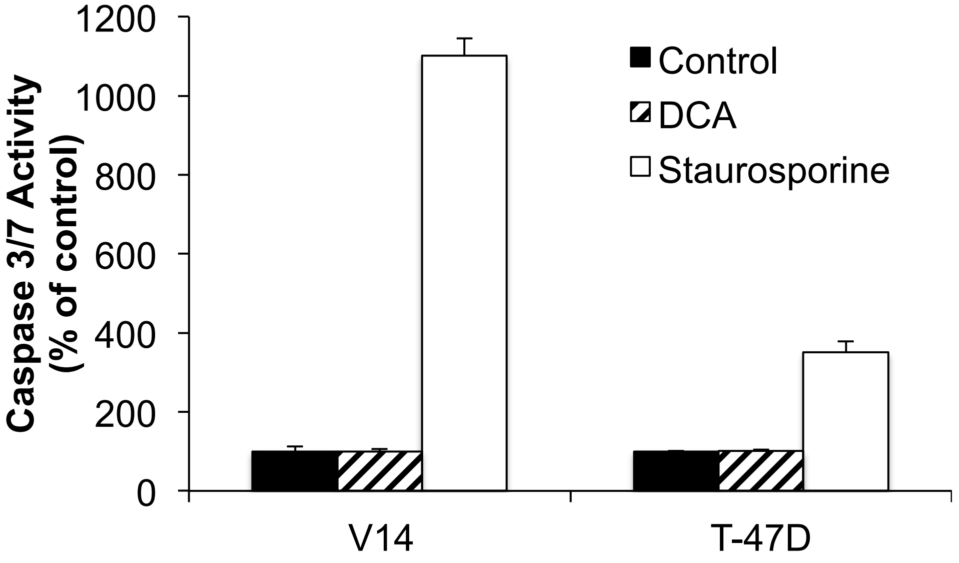


**Supplementary Figure 3**

Caspase 3/7 activity in V14 and T-47D cell lines after 12 hr treatment with DCA (5mM) or staurosporine (1 uM, positive control for apoptosis).

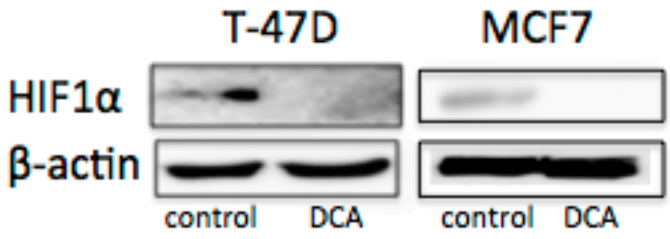


**Supplementary Figure 4**

HIF1$\alpha$ protein expression by western blotting after treatment with 1 mM DCA for 72 hrs.
