## Supplementary Methods for "Pyruvate dehydrogenase kinase expression profile is a biomarker for cancer sensitivity to dichloroacetate-mediated growth inhibition"

**Supplementary Information**

### Methods

#### Cell culture

Human cancer cell lines were obtained from (year obtained in brackets): Breast cancer cell lines: Dr Anna DeFazio, Westmead Millenium Institute, Sydney, Australia provided T-47D (2003), MCF-10A (2005, non-cancerous); Prof Chris Parish, Australian National University, Canberra, Australia provided MDA-MB-468 (2003) MCF7 (2003) and MDA-MB-231 (2003); June Hornby, Australian National University, provided PC-3 prostate cancer (2000), SW620 colon cancer (2000); Dr Pierre Dilda Lowy Cancer Center, UNSW, Australia provided Bx-PC3 pancreatic cancer (2011). 4T1 mouse mammary tumor cells were obtained from Prof Robin Anderson, Peter MacCallum Cancer Centre, Melbourne, Australia (2007). The cell lines have appearances consistent with published morphologies. MCF10A cells were grown in DMEM/F-12 medium, 25% horse serum, 0.01% EGF, 0.28 IU/ml insulin, 0.01% cholera toxin and 0.5µg/ml hydrocortisone.

#### V14 mouse mammary tumor cell line

The V14 cancer cell line was derived by Dr. Anneke Blackburn from a spontaneous mammary adenocarcinoma arising at 40 weeks of age in a BALB/c *Trp53*$^{+/-}$ mouse (Blackburn *et al*, 2004). The original tumor and transplanted outgrowths from this tumor have previously been determined to be estrogen receptor and progesterone receptor negative, HER2/neu positive, and to have lost the wild-type allele of *Trp53*. This tumor line was not metastatic (Yan *et al*, 2010). V14 cells were grown in DMEM/F-12 medium supplemented with 25 mM Hepes (MP Biomedicals, USA), 1.2 g/L $NaHCO_3$, 2% adult bovine serum (ABS), 0.1% PSN (3% penicillin, 5% streptomycin and 5% neomycin), 5 ng/ml epidermal growth factor (EGF), 10 µg/ml insulin, 15 µg/ml gentamicin (Sigma, MO, USA).

**In vivo tumor treatment with DCA**

Animal experiments were conducted with the approval of the Australian National University Animal Ethics Experimentation Committee under the guidelines established by the Australian National Health and Medical Research Committee. Female BALB/c mice, 8-14 weeks old, were anaesthetized with isofluorane and injected subcutaneously on the flank with prepared tumor cells in 15 µl of serum-free media. For V14 tumors, 5 x $10^6$ cells were injected in each the left and right flank. Tumors were allowed to become established to approx. 50 $mm^3$ size before commencing treatment with DCA on day 11. For 4T1 tumors, 1 x $10^5$ cells were injected in one site per mouse. As 4T1 cells were considerably less sensitive to DCA in vitro than V14 cells, treatment of the mice bearing 4T1 tumors was commenced 3 days prior to injection of the 4T1 cells to maximize the delivery of DCA to the tumors. (In vivo, DCA inhibits its own metabolism via the irreversible inactivation of GSTZ1 (Anderson *et al*, 1999), thus the bioavailability of DCA increases with treatment time.) DCA was administered in the drinking water at 1.5 g/L, and water consumption was monitored by weighing the water bottles every 2-4 days. The DCA did not alter water consumption significantly, and delivered approximately 115 or 160 mg/kg/day for the 4T1 and V14 experiments respectively. The early development of 4T1 tumors was not affected by DCA pre-treatment. Tumor size was monitored every 2-3 days with electronic calipers and the tumor size estimated using the formula $\pi/6*l*w^2$, where l is the longest dimension, and w is the shorter dimension perpendicular to l. Upon sacrifice, tumor tissues were fixed overnight in neutral buffered formalin, and processed for H&E staining. The number of mitotic figures present in H&E sections was counted per 10 high powered fields (hpf) per tumor, with the investigator blind to the treatment groups.
